# Dopaminergic signaling supports auditory social learning

**DOI:** 10.1101/2021.03.03.433719

**Authors:** Nihaad Paraouty, Catherine R. Rizzuto, Dan H. Sanes

## Abstract

Explicit rewards are commonly used to reinforce a behavior, a form of learning that engages the dopaminergic neuromodulatory system. In contrast, skill acquisition can display dramatic improvements from a social learning experience, even though the observer receives no explicit reward. Here, we test whether a dopaminergic signal contributes to social learning in naïve gerbils that are exposed to, and learn from, a skilled demonstrator performing an auditory discrimination task. Following five exposure sessions, naïve observer gerbils were allowed to practice the auditory task, and their performance was assessed across days. We first tested the effect of an explicit food reward in the observer’s compartment that was yoked to the demonstrator’s performance during exposure sessions. Naïve observer gerbils with the yoked reward learned the discrimination task significantly faster, as compared to unrewarded observers. The effect of this explicit reward was abolished by administration of a D1/D5 dopamine receptor antagonist during the exposure sessions. Similarly, the D1/D5 antagonist reduced the rate of learning in unrewarded observers. To test whether a dopaminergic signal was sufficient to enhance social learning, we administered a D1/D5 receptor agonist during the exposure sessions in which no reward was present, and found that the rate of learning occurred significantly faster. Finally, a quantitative analysis of observer vocalizations and movements during the exposure sessions suggest behavioral strategies that contribute to social learning. Together, these results are consistent with a dopamine-dependent reward signal during social learning.

## Introduction

A broad range of behavioral paradigms demonstrate that *explicit* rewards, such as food or money, can reinforce behaviors and facilitate learning (Hull, 1943; Ferster, 1957; Tempel et al., 1983; Schultz, 2000; O’Doherty, 2004; Nakatani et al., 2009; Gottlieb et al., 2014; Berridge, 2007; Robbins and Everitt, 2007; Steinberg et al., 2013). These reinforcing rewards typically engage dopaminergic signaling which acts to modulate both motivation and memory formation (Wickens et al., 2003; Schultz, 2002; Lisman and Grace, 2005; Calabresi et al., 2007; Rossato et al., 2009; Bromberg-Martin et al., 2010; Flagel et al., 2011). Specifically, dopamine neuron activity in pars compacta of substantia nigra and the ventral tegmental area is correlated with reward presentation, as well as reward anticipation (Schultz et al., 1997; Schultz, 2013). Furthermore, anticipatory dopaminergic activity is thought to guide and accelerate learning through D1 dopamine receptor-dependent long-term potentiation (Schultz, 2006; Calabresi et al., 2007; Adamantidis et al., 2011). Reward anticipation can also enhance attention and, in turn, boost the encoding of incoming sensory signals (Maunsell et al., 2004; Seitz and Watanabe, 2005; Sasaki et al., 2010). Thus, explicit rewards have a positive impact on the learning and retention of sensory and motor skills (Roelfsema et al., 2010; Wickens et al., 2003; Abe et al., 2011; Orban et al., 2011; Shmuelof et al., 2012; Galea et al., 2015).

The role of dopaminergic signaling in explicit reward learning suggests that dopamine may also be engaged by *implicit* rewards, such as those triggered by learning in the absence of any external feedback (Ripolles et al., 2016; 2018), or certain intrinsic motivational states such as curiosity (Gruber et al., 2014). In principle, implicit reward signals share some of the neural mechanisms that attend the acquisition of an external reward, including dopamine release in the nucleus accumbens (Radhakishun et al., 1988; Hernandes and Hoebel, 1990; Martel and Fantino, 1993; Roitman et al., 2004). For example, fluctuations in dopamine levels occur in rat nucleus accumbens when naïve animals observe the delivery of an explicit reward to a conspecific (Kashtelyan et al., 2014). In these experiments, the naïve observer rat experienced an initial increase in dopamine levels, followed by a decrease in the absence of an explicit reward. Similarly, Ripolles et al. (2014; 2016) showed that when subjects successfully learn the meaning of new words presented in verbal contexts, in the absence of any explicit reward or feedback, they also experienced an increase in emotion-related physiological measures and subjective pleasantness ratings. Such learning in the absence of external reward or feedback was found to be causally related to synaptic dopamine availability (Ripolles et al., 2018). When the subjects were provided with a dopamine precursor or a dopamine antagonist, both learning and pleasantness ratings were shifted as compared to a placebo group. The precursor group showed enhanced learning and pleasantness ratings, while the antagonist group showed a decrease in learning and pleasantness ratings.

Social interactions, themselves, can act as rewards (Panksepp and Lahvis, 2017; Dölen et al., 2013; Hung et al., 2017). Access to social stimuli or social isolation both recruit the dopamine reward system (Gunaydin et al., 2014; Matthews et al., 2016; Tamir and Hughes, 2018). In the current study, we asked whether dopamine signaling during exposure to a performing conspecific could contribute to social learning, defined here as the acquisition or facilitation of new skills by observation or exposure to a conspecific performing a well-defined behavior (Carcea and Froemke, 2019; Paraouty et al., 2020). We previously reported that naïve gerbils acquire a sound discrimination task significantly faster when exposed for five days to a demonstrator that was performing the task, as compared to three different control groups (Paraouty et al., 2020). To test the hypothesis that social learning engages a dopamine-dependent reward signal that facilitates the subsequent acquisition of an auditory task, we used pharmacological loss- and gain-of-function manipulations of D1/D5 dopamine receptors in naïve observer gerbils during exposure sessions. Furthermore, to assess the behavioral signals that might contribute to social learning, we monitored the vocalizations and movements of both observer and demonstrator during the exposure sessions. Together, the results suggest that dopaminergic signaling is both necessary and sufficient to facilitate socially-mediated task acquisition, and one important social cue is the demonstrators’ vocalizations at trial initiation.

## Results

### Presence of an explicit reward improves social learning

Our first objective was to determine whether an explicit reward could facilitate social learning. As described previously (Paraouty et al., 2020), demonstrator gerbils were first trained by an experimenter to perform a Go-Nogo amplitude modulation (AM) rate discrimination task. Briefly, the demonstrators were placed on controlled food access and trained to initiate each trial by placing their nose in a nose port. The Go stimulus (12-Hz AM noise) indicated the presence of a food reward at the food tray, while the Nogo stimulus (4-Hz AM noise) signaled the absence of a food reward. A discrimination performance metric, d-prime (d’) was calculated for each session as d’ = z(hit rate) – z(false alarm rate). Once animals performed the task with a d’ > 1.5, they qualified as a demonstrator (for more details, see Methods Section).

Our previous results demonstrated that observation alone led to social learning (Paraouty et al., 2020), and we first replicated that finding. A naïve observer gerbil was placed adjacent to a previously-trained and performing demonstrator gerbil for 5 consecutive exposure sessions (Figure 1A, left, also see Video 1). The demonstrator compartment was separated from the observer compartment by a transparent divider, and possessed a nose port and a food tray, thereby allowing the demonstrator gerbil to initiate and perform trials. The naïve observer gerbil had access to all sensory cues emanating from the demonstrator, and to the Go and Nogo sounds delivered 1 m above the test cage. During each exposure session, the naïve observer gerbil was exposed to a minimum of 80 Go trials and 20 Nogo trials performed by the demonstrator (Figure 1B, brown lines). Following the fifth exposure session, the demonstrator was removed as well as the divider, and the naïve observer gerbil was permitted to practice the task on its own (Figure 1A, right). The sensitivity metric, d’, was computed for all sessions during which the naïve observer gerbil performed > 15 Nogo trials. On average, the naïve observer gerbils required 5.6 ± 0.35 days (mean ± standard error) to perform the task at a criterion d’ of 1.5 (Figure 1C, black lines). No significant difference was found between the number of days taken to reach the criterion d’ by the current observers and those tested in Paraouty et al. (2020, Figure 1, Wilcoxon rank sum test, X^2^(1)=0.27, *p*=0.601). Therefore, the current study contains a replication of our previous finding that social learning facilitates acquisition of a sound discrimination task (Paraouty et al., 2020) as compared to 3 control conditions that were examined in the original study: 1) naïve gerbils deprived of any prior exposure before performing the task, 2) naïve gerbils exposed to the test cage alone for 5 days prior to performing the task, and 3) naïve gerbils exposed to a non-performing demonstrator for 5 days prior to performing the task. The 3 control groups all took significantly longer to perform the task and reach the criterion d’ of 1.5 as compared to the observer groups (see Figure 2 in Paraouty et al., 2020).

**Figure 1.**
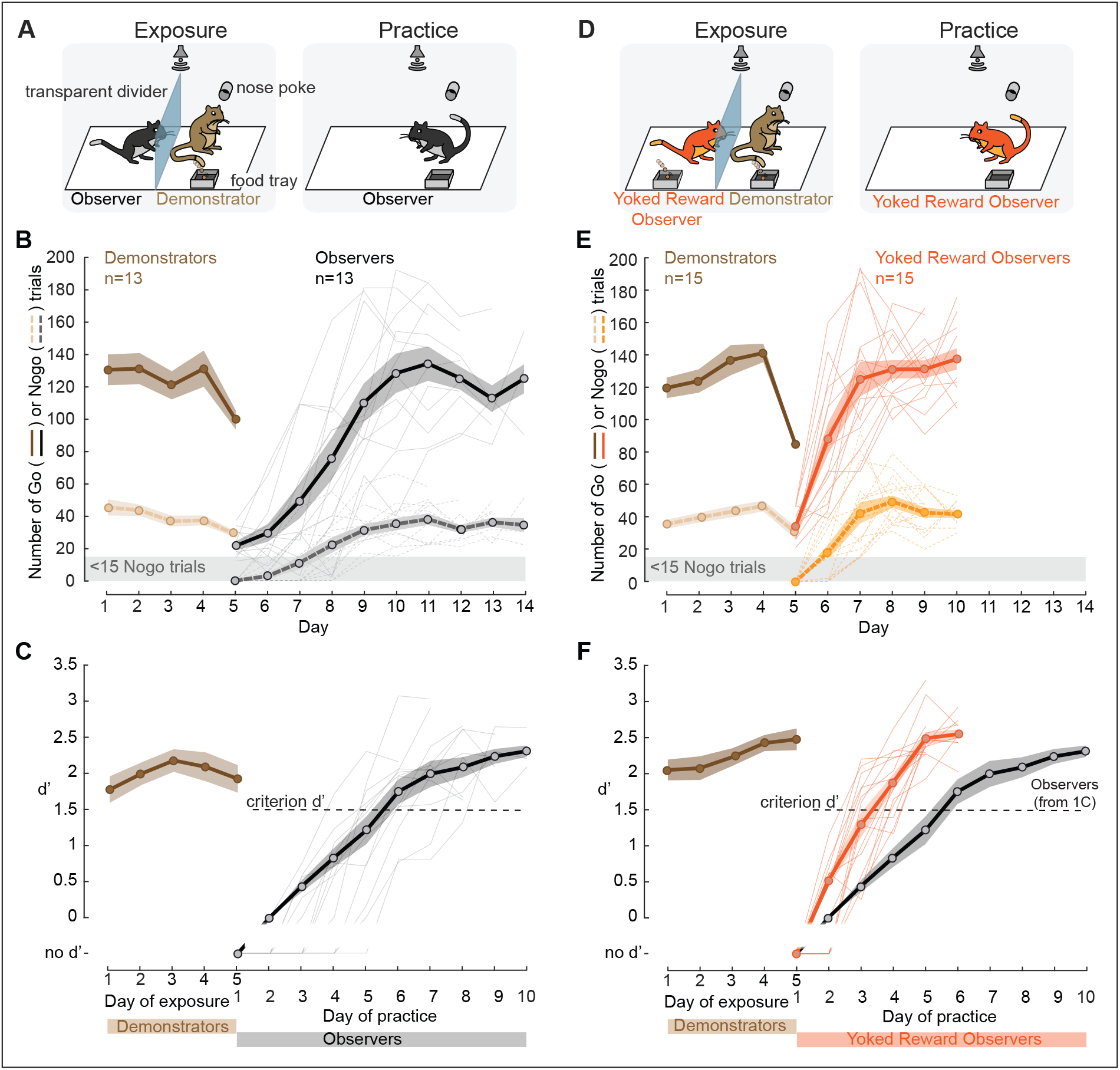
Yoking food reward to an observer facilitates social learning. **A.** Replication of Figure 1 from Paraouty et al. (2020). Experimental Design. Left: Naïve observer gerbil (black) was separated from a performing same-sex demonstrator gerbil (brown) by a transparent divider. The demonstrator was previously trained by the experimenters (see Methods for details). Right: Following five days of exposure, the practice phase began during which the naïve observer gerbil was permitted to practice the task on its own. **B.** Individual (thin lines) and mean (thick lines) ± standard error of the mean (SEM) of overall number of Go (dark color) and Nogo trials (pale color) performed by the demonstrators (5 males; age=122.6±4.7 days (mean±standard error)) during the 5 days of exposure (brown), and by the naïve observer gerbils (age=101.4±4.1) during the practice sessions (black). **C.** Individual (thin lines) and mean ± SEM (thick lines) performance d’ values of the demonstrator during the 5 days of exposure (brown) and of the naïve observer gerbil during the practice sessions (black). The performance sensitivity, d’ was calculated once the observers performed >15 Nogo trials. **D.** Experimental Design. Left: Yoked reward observer gerbil (orange) was separated from a performing demonstrator gerbil (brown) by a transparent divider. A food tray was also present in the observer’s compartment, and the food reward of the demonstrator animal was yoked to that of the observer. Right: Practice session, similar to A (right). **E**. similar to B, with demonstrators (4 males; age=130±4.2) and yoked reward observers (age=96.8±5.3). **F.** similar to C for the yoked reward observers. The pale grey line indicates the mean d’ of the naïve observers without the yoked reward (from Figure 1C).

**Figure 2.**
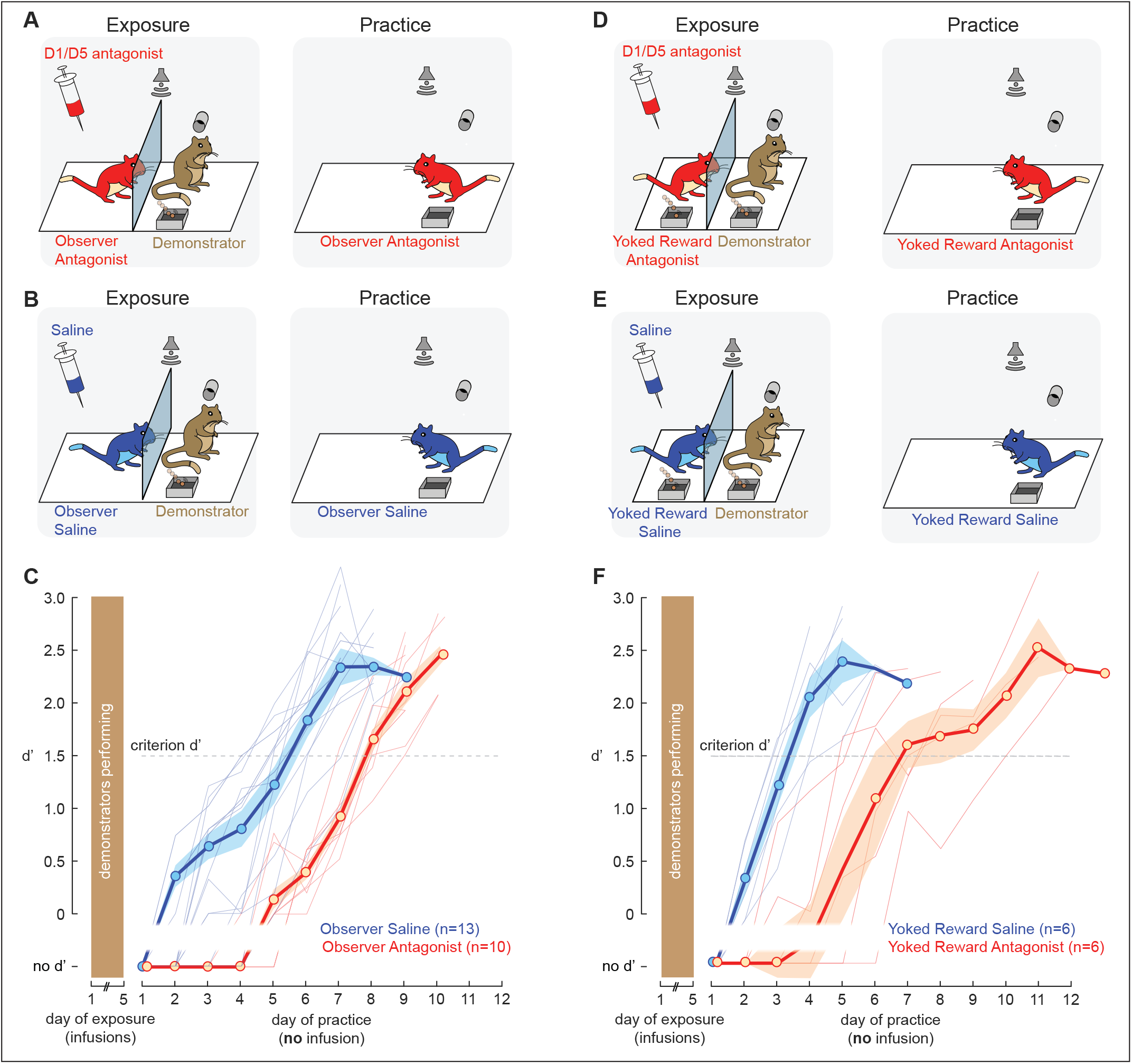
Decrease of tonic dopamine level degrades social learning. **A.** Prior to each exposure session, the naïve observer gerbils were given an injection of a dopamine receptor antagonist (5 males; age=99.5±8.6 days; see Methods for details). After 15-20 min of recovery, the exposure session began (left) with same-sex demonstrators (age=138±3.7). No drug was injected during the practice sessions (right). **B.** Similar to A, except that saline was injected to the observers (5 males; age=104.3±10.7) prior to each exposure session (left) with the demonstrators (age=143±5.4). No drug was injected during the practice sessions (right). **C.** The performance of the observer gerbils which received a dopamine antagonist (red) and those that received saline (blue) as a function of practice sessions. **D.** Prior to each exposure session, the yoked reward observer gerbils were given an injection of a dopamine receptor antagonist (4 males; age=111.3±8.9). After 15-20 min of recovery, the exposure session began (left) with the demonstrators (age=166±4.5). No drug was injected during the practice sessions (right). **E.** Similar to D, except that saline was injected to the observers (5 males; age=110.3±8.5) prior to each exposure session (left) with the demonstrators (age=119±5.7). No drug was injected during the practice sessions (right). **F.** The performance of the yoked reward observer gerbils which received a dopamine antagonist (red) and those that received saline (blue) as a function of practice sessions.

To test whether an explicit food reward could further enhance social learning, we next placed a food dispenser in the observation chamber (Figure 1D, left), and provided naïve observers with a food reward that was yoked to the demonstrator’s performance during the five exposure sessions. In other words, a food pellet was delivered to both the demonstrator and the naïve observer when the demonstrator responded accurately on Go trials. The naïve observers in this case were also exposed to a minimum of 80 Go trials and 20 Nogo trials in each pairing session (Figure 1E, brown lines). On average, the naïve observer gerbils with an explicit yoked reward required 3.3 ± 0.21 days to perform at criterion (Figure 1F, orange lines) which was significantly faster, as compared to observation alone (in grey; Wilcoxon rank sum test, X^2^(1)=12.87, *p*=0.0003). Therefore, an explicit food reward during the exposure sessions facilitates social learning.

### Decreasing dopamine receptor signaling diminishes social learning

Next, we asked whether an intrinsic dopaminergic signal during the exposure sessions contributed to social learning. Prior to each exposure session, the naïve observer gerbils were briefly anaesthetized with isoflurane and given an intra-peritoneal injection of a D1/D5 dopamine receptor antagonist, SCH-23390 (0.03 mg/kg, Figure 2A, left) or an injection of saline (Figure 2B, left). Animals were allowed to recover for 15-20 minutes before the exposure session began. Following the five days of exposure, the naïve observer animals were allowed to practice the task, and no further injections were delivered (Figure 2A-B, right). When D1/D5 receptor were blocked during the exposure sessions, naïve observer gerbils took significantly longer to reach the criterion performance, as compared to the saline-injected observer animals (Figure 2C; Wilcoxon rank sum test, X^2^(1)=13.81, *p*=0.0002). The saline-injected animals did not differ significantly from the uninjected animals (from Figure 1C; Holm-Bonferroni-corrected post-hoc comparisons, *p*=0.7805), suggesting that neither saline nor the handling of the animal for the injection influenced the rate of task acquisition. During the first practice sessions, both trial initiation and hit rates were significantly poorer for the antagonist-injected animals as compared to all other groups (see Supplementary Figure 2, Holm-Bonferroni-corrected post-hoc comparisons, *p*<0.05), suggesting that the five exposure sessions did not benefit those animals. Thus, a decrease in tonic dopamine level during the exposure sessions led to a significant decrease in the subsequent rate of task acquisition.

We also tested whether a decrease in an intrinsic dopaminergic signal could be compensated for by the presence of an external food reward during the exposure sessions. Two additional groups of naïve observer gerbils were briefly anaesthetized prior to each exposure session, and given an intra-peritoneal injection of a D1/D5 dopamine receptor antagonist, SCH-23390 (0.03 mg/kg, Figure 2D, left) or an injection of saline (Figure 2E, left). After recovery, both groups of naïve animals were provided with the exposure sessions, during which they received a food reward that was yoked to the demonstrator’s performance. Following the five days of exposure, the yoked reward observer animals were allowed to practice the task, and no further injections were delivered (Figure 2D-E, right). Naïve observer gerbils took significantly longer to reach the criterion performance when D1/D5 receptor were blocked, despite the presence of the yoked food reward during exposure sessions, as compared to the saline-injected control group (Figure 2F; Wilcoxon rank sum test, X^2^(1)=8.67, *p*=0.0032). In addition, the saline-injected animals did not differ significantly from the uninjected animals with the yoked reward (from Figure 1F; Holm-Bonferroni-corrected post-hoc comparisons, *p*=0.8861), suggesting that neither saline nor the handling of the animal for the injection influenced the rate of task acquisition. Trial initiation and hit rates were significantly impacted during the first practice sessions for the antagonist-injected animals despite the presence of the yoked reward during the exposure sessions (see Supplementary Figure 2, Holm-Bonferroni-corrected post-hoc comparisons, *p*<0.05). Thus, these results confirm that a decrease in intrinsic tonic dopamine level during the exposure sessions lead to a significant decrease in the subsequent rate of task acquisition despite the presence of yoked food reward. Furthermore, comparison of both observers groups which received the dopamine antagonist revealed that observers with the yoked food reward did not perform significantly better than the one without the yoked food reward (Holm-Bonferroni-corrected post-hoc comparisons, *p*=0.0807). Together, these results suggest a crucial role of intrinsic dopamine signaling in social learning.

### Pharmacological increase in tonic dopamine level facilitates social learning rate

In order to test whether a high intrinsic dopamine level was enough to facilitate social learning, naïve observer animals received a D1/D5 dopamine receptor agonist during the exposure sessions. Prior to each exposure session, naïve observer gerbils were briefly anaesthetized and given either an intra-peritoneal injection of a D1/D5 dopamine receptor agonist, SFK-38393 (5.0 mg/kg, Figure 3A, left) or an injection of saline (Figure 3B, left). Animals were allowed to recover for 15-20 minutes before the exposure session began. Following the five days of exposure, the animals were allowed to practice the task, and no further injections were delivered (Figure 3A-B, right). When D1/D5 receptors were activated during exposure sessions, naïve observer gerbils subsequently learned the task significantly faster, as compared to a group of saline-injected animals (Figure 3C; Wilcoxon rank sum test, X^2^(1)=9.84, *p*=0.0017). Thus, an increase in intrinsic dopamine signaling during the exposure sessions led to enhanced task acquisition during the subsequent practice sessions.

**Figure 3.**
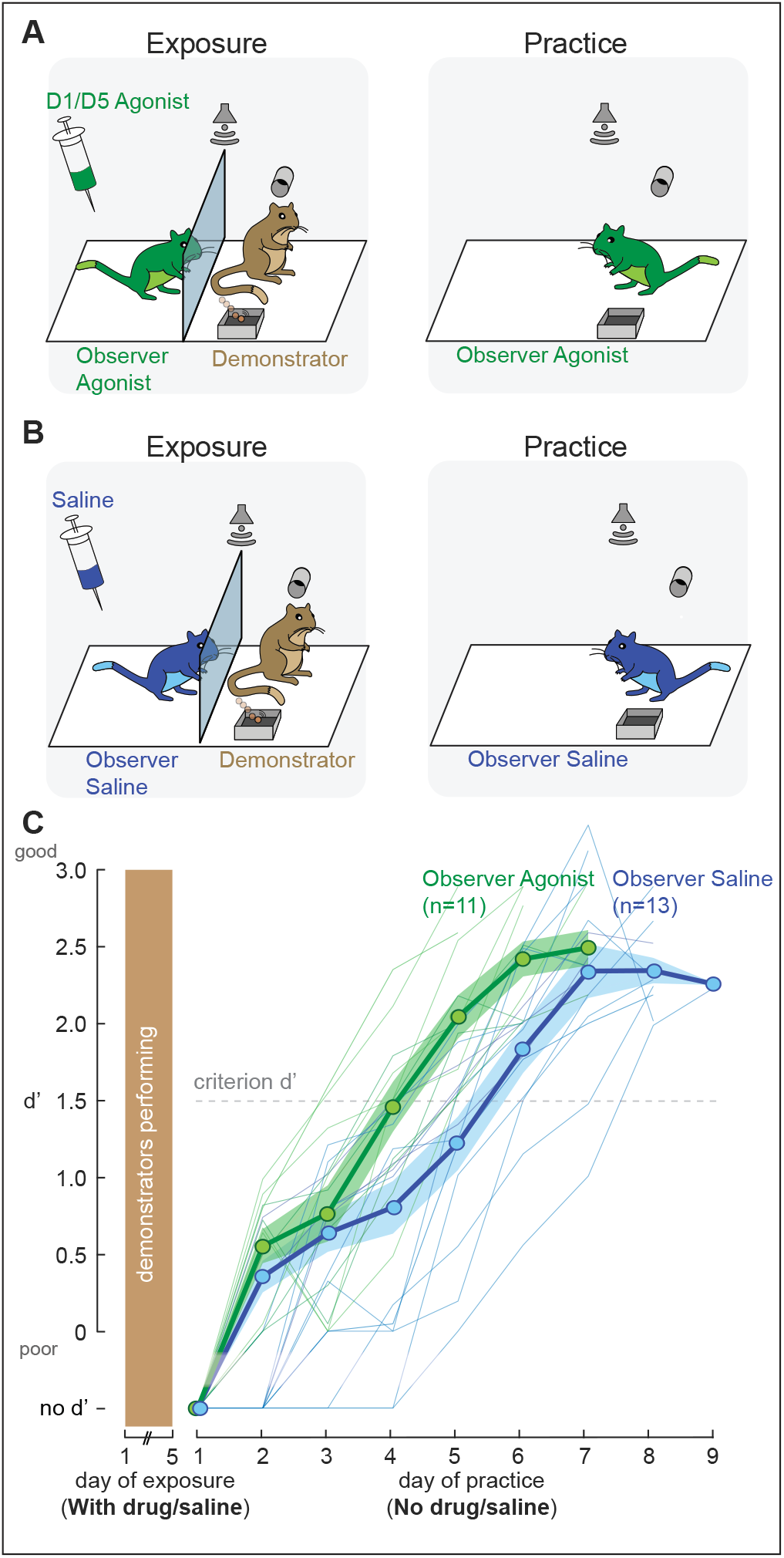
Increase of tonic dopamine level enhances social learning. **A.** Prior to each exposure session, the naïve observer gerbils were given an injection of a dopamine receptor agonist (6 males; age=100.3±12 days; see Methods for details). After 15-20 min of recovery, the exposure session began (left) with the demonstrators (age=139±5.2). No drug was injected during the practice sessions (right). **B.** Similar to A, except that saline was injected to the observers (5 males; age=104.3±10.7) prior to each exposure session (left) with the demonstrators (age=163±5.4). No drug was injected during the practice sessions (right). **C.** The performance of the observer gerbils which received a dopamine agonist (green) and those that received saline (blue) as a function of practice sessions.

### D1/D5 antagonist during exposure changes the behavioral pattern of the observer animals

Video footage from the five daily exposure sessions was recorded (see Video 1) and analyzed for a subset of observer animals (n=5) in each of the seven experimental groups tested. To standardize the analyses, we only included the frames obtained during the first 20 minutes of each exposure session. Both the observer animal’s position (Supplementary Figure 1A) and its head orientation (Supplementary Figure 1C) were computed using DeepLabCut (Mathis et al., 2018, see Methods). On average, no significant difference was found between observer animals with or without a yoked reward (Supplementary Figure 1B, left, Holm-Bonferroni-corrected post-hoc comparisons, *p*=0.7859) in terms of distance travelled during each exposure session. However, observer animals which received the dopamine antagonist moved significantly less as compared to observer animals which received either saline or a dopamine agonist (Supplementary Figure 1B, middle, *p*=0.0192 and *p*=0.0012). In contrast, observers which received the dopamine agonist did not differ significantly from the saline observer animals (*p*=0.3424). Yoked reward observers with the dopamine antagonist also moved significantly less as compared to the yoked reward saline controls (Supplementary Figure 1B, right, *p*=0.015).

The head orientation of all observer groups were preferentially towards the observer’s food tray, whether or not a food tray was present in the observer’s compartment during the exposure sessions. However, in the presence of the food tray, the yoked reward observers oriented their head significantly more towards the food tray as compared to observers without a food tray (Supplementary Figure 1D, left, *p*=0.0011). Interestingly, observers with the dopamine antagonist which moved significantly less, also showed less preference to the food tray location as compared to observers with saline or with dopamine agonist (Supplementary Figure 1D, middle, , *p*<0.0001 for both). Similarly, the yoked reward observers which received the dopamine antagonist showed a significantly reduced preference to the food tray as compared to the saline controls (Supplementary Figure 1D, right, *p*=0.0019). Although the dopamine antagonist impacted both locomotion and head orientation, the yoked reward observers with dopamine antagonist consumed a similar number of food pellets during the exposure sessions as the yoked reward observers with saline or the yoked reward uninjected observers (*p*=0.776; p=0.053). In addition, for all observers in the yoked reward condition (uninjected, saline-injected, and antagonist-injected observers), no correlation was found between the mean number of pellets consumed by the observers during the five exposure days and the subsequent number of practice days taken to reach the criterion d’ (r=0.17; *p*=0.399). Together, these results confirm that the trend towards reduced food intake of the antagonist-injected observers as compared to the uninjected animals during the exposure sessions was not a limiting factor for the subsequent learning delay during the practice sessions.

### The timing of vocalizations during exposure matches closely the onset of the sound stimuli

Audio recordings from the five daily exposure sessions were obtained and analyzed using DeepSqueak (Coffey et al., 2019, see Methods) for the same subset of animals used for video analyses (n=5 in each of the seven observer groups tested). To standardize the analyses, we only included calls that were present during the first 20 minutes of each of the exposure session. Interestingly, both demonstrator and observer animals vocalized during the exposure sessions (Figure 4A). In general, male observers displayed significantly longer call durations and lower principal frequencies, as compared to female observers (see Figure 4B; Steel-Dwass comparison, *p*< 0.0001 for both vocalization parameters). The two groups of observers which received the dopamine antagonist vocalized significantly less, as compared to their respective saline controls (Figure 4C, observers in the unyoked reward condition, *p*=0.0006; observers in the yoked reward condition, *p*=0.0011).

**Figure 4.**
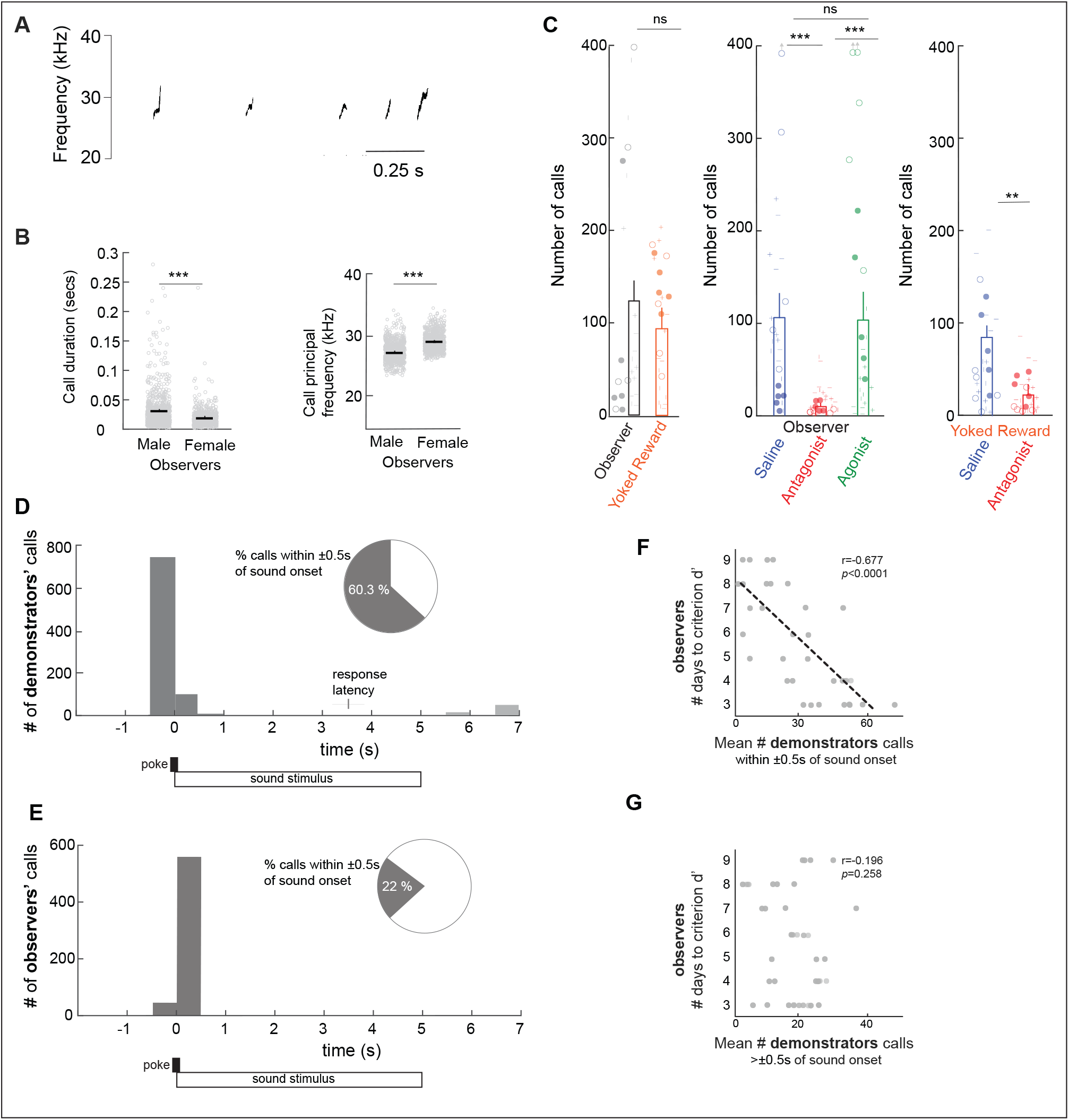
Synchrony between vocalization and sound onset may provide key social cues to observers. **A.** Example of a series of vocalizations of a naïve observer during an exposure session. **B.** Sex differences in terms of call duration (left) and call frequency (right) for all pairs of observer and demonstrator. **C.** Mean number of calls of 5 observers (shown in different symbols) in each of the seven observer groups during the first 20 minutes of the audio recordings for each of the five daily exposure sessions. **D.** Histogram of vocalization onsets for a subset of 5 demonstrators in each experimental condition as a function of the onset of the Go and Nogo sound stimuli. The inset represents the percent of demonstrators’ calls that were present within ±0.5s of the sound onset (dark grey bars). **E.** Histogram of vocalization onsets for a subset of 5 observers in each experimental condition as a function of the onset of the Go and Nogo sound stimuli (the same demonstrator-observer pairs recordings were analyzed). The inset represents the percent of observers’ calls that were present within ±0.5s of the sound onset (dark grey bars). **F.** Mean number of demonstrators’ calls within ±0.5s of the sound onset plotted against the number of days taken by the respective observers to reach the criterion d’ of 1.5. The data is from 5 demonstrator and observer pairs in each of the seven experimental groups tested (average of the five daily exposure sessions). **G.** Mean number of demonstrators’ calls unsynchronized to the sound onset plotted against the number of days taken by the respective observers to reach the criterion d’. Asterisks denote statistically significant post-hoc differences at the following levels: * *p*<0.05 and \*\**p*<0.01.

For all exposure sessions analyzed, 60% of the demonstrators’ vocalizations were initiated around the time of trial initiation (i.e., nose-poke) and presentation of the Go or Nogo sound stimulus (Figure 4D, dark grey bars). While a similar temporal relationship was displayed by the naïve observers (Figure 4E), these animals produced fewer vocalizations around the presentation of the Go or Nogo sound stimulus (22% of all calls). Furthermore, while a majority of those demonstrators’ calls occurred prior to the nose poke and sound onset, the majority of the observers’ calls occurred after the sound onset (Figures 4D and 4E, respectively).

To further assess whether the timing of the demonstrators’ vocalizations influenced the observers’ rate of social learning, we asked whether there was a relationship between the proportion of demonstrators’ calls that occurred within ±0.5s of the sound onset during the exposure sessions, and the observers’ subsequent rate of task acquisition during the practice sessions. In fact, there was a strong positive correlation (Figure 4F; Pearson’s correlation coefficient, r=−0.677, *p*< 0.001). In contrast, no correlation was found between the percent of demonstrator calls that occurred outside of the sound onset (i.e., >0.5s before or after the Go/Nogo stimulus) and the subsequent learning rate of the observers (Figure 4G; r=−0.197, *p*=0.258). Overall, these results suggest that the timing of the demonstrators’ vocalizations with the sound onset during the exposure sessions is a possible social cue for the observers to learn in this particular context.

### Non-task factors do not account for differences in learning rates

A range of experimental parameters could have contributed to the differences in learning rates observed across the different groups tested, including sex and postnatal age. For all groups tested, both male and female observers were used. No significant difference was found between male and female gerbils in terms of number of days to reach a criterion d’ of 1.5, when combined across groups (Kruskal-Wallis H test, X^2^(6)= 3.37, *p*=0.770) or within each group (*p*>0.05). The mean postnatal age of the different observer groups did not differ significantly (Kruskal-Wallis H test, X^2^(6)= 6.51, *p*=0.369). No significant group difference was found in terms of total number of trials performed and d’ measures of the different demonstrator groups during the 5 exposure sessions (Kruskal-Wallis H test, X^2^(6)= 4.84, *p*=0.564; X^2^(6)= 6.1, *p*=0.423, respectively). Together, these results suggest that the differences in social learning rates could not be explained by non-task factors.

## Discussion

Dopamine signaling plays a broad role in learning that results in acquisition of explicit rewards (review, Schultz, 2016), suggesting that it may play a general role in social forms of learning that do not yield an immediate reward. To test this idea, dopamine signaling was manipulated only during the periods of social exposure. We first established that an explicit food reward during social exposure could facilitate the subsequent rate of learning (Figure 1). We then tested whether a reduction in tonic dopamine signaling caused a reduction in social learning. We found that the rate of learning was significantly delayed when animals were treated with a D1 receptor antagonist during social exposure (Figure 2). Finally, to test whether tonic dopamine signaling was sufficient to facilitate social learning, in the absence of any external reward, animals were treated with a D1 receptor agonist during social exposure (Figure 3). This lead to faster task learning, suggesting a role for dopaminergic signaling during social learning itself, which is distinct from any dopaminergic signaling that may occur while animals practice the task and begin to receive an explicit food reward. Overall, our findings suggest that theories of dopamine-dependent learning can account for task acquisition paradigms in which explicit rewards, and reward prediction errors, are unavailable.

Our results are broadly consistent with the role of dopaminergic signaling during social interactions. Indeed, dopamine has been found to play a vital role in social transmission of food preferences (Matta et al., 2017; Rodriguiz et al., 2004). For example, social learning of food preference is impaired when observer mice receive a D1-type receptor antagonist, whereas a D2 receptor antagonist is ineffective (Choleris et al., 2011). Similarly, dopaminergic signaling is involved in learned aggression in rats (Suzuki and Lucas, 2015). Although the present study was not designed to assess the role of D1/D5 signaling on attention, it is plausible that the agonist led to enhanced arousal and attention during the exposure sessions (Bellgrove and Mattingley, 2008; Noudoost and Moore, 2011; Thiele and Bellgrove, 2018). In contrast, the antagonist may have decreased the general arousal and attention of the observers to both the social and non-social cues (to the demonstrator animal and the sound stimuli, respectively), or led to an aversive response to the test cage (Acquas and Di Chiara, 1994) in addition to the decrease in internal dopamine signaling.

In our experiments, tonic dopamine levels in the naïve observers were modulated during each exposure session through systemic drug injections. However, dopamine signaling is often temporally precise at the moment when animals receive a reward or perceive a cue that predicts a reward (Schultz, 2000). Therefore, it is possible that our manipulations also enhanced or depressed the transient dopamine signals occurring during the exposure sessions. For example, increased firing of dopamine neurons in the observers could have been elicited by specific actions of the demonstrator, such as nose poking, or food acquisition, or vocalization. The yoked-reward observers in our study generally did not seek, or consume, the food pellet immediately after its delivery during the exposure sessions. Therefore, it is plausible that both a general increase in dopaminergic activity, as well as a temporally precise signal (demonstrators’ vocalizations) could have elicited dopaminergic activity transients, and each served to facilitate an expectation for sound cues and potential rewards. Future work would require that dopamine neuron activity is monitored during the observation period to directly determine whether signaling is tonic or phasic.

Auditory forms of social learning have been particularly well-studied in songbirds (Mooney, 2009; Chen et al., 2016; Mennill et al., 2018; Narula et al., 2018 ; Yanagihara & Yazaki-Sugiyama, 2019), where dopaminergic signaling is involved in both song learning and song production (Kubikova et al., 2009; Gale and Perkel, 2010). For example, heightened dopamine signaling was shown in Area X during directed singing, and this may facilitate learning (Sasaki et al., 2006). Similarly, local stimulation of D1/D5 receptors in the auditory cortex has been found to boost auditory stimulus detection (Bao et al., 2001; Happel et al., 2014; Deliano et al., 2020).

As shown in Paraouty et al (2020), gerbils are able to use either visual or auditory cues during the exposure period. Furthermore, they have access to olfactory cues which may heighten arousal, and somatosensory cues which may permit adaptation to the general testing environment. Therefore, we suggest that observer animals are using these modalities, either individually or in combination during the course of a single exposure session, and learning would accrue from each. One intriguing possibility is that observers attend to the demonstrators’ vocalizations which were closely linked to a behavior (nose poke) and the sound stimulus onset (Figure 4). We speculate that the timing of the demonstrators’ calls could boost the observers’ attention to both the nose-poke behavior and the sound stimuli, thereby promoting the formation of visual memories of the demonstrator’s position near the poke, or auditory memories of a sound cue associated with a vocalization, or both. While demonstrators’ vocalizations may convey crucial social cues to the observers, when matched with the onset of the auditory Go or Nogo stimuli, future experiments could assess the individual role of each social cue.

In the present study, the task to be learned through social exposure involved several discrete behaviors: trial initiation by nose poking, reward-seeking following a Go stimulus, and the withholding of reward-seeking following a Nogo stimulus. The movements of the naïve observer gerbils during the exposure sessions indicate some nascent forms of imitation, with a preferred head direction towards the food tray location (Supplementary Figure 1C-D), suggesting one element of the task that may be learned early on, even in the absence of visual cues (Ellard and Eller, 2009). This head orientation preference may be explained, at least in part, with (1) the cage design that was less wide than long, (2) the lateral position of rodent eyes which permit attention while at a right angle to visual objects, and (3) the gerbil’s ability to attend to auditory cues (Paraouty et al., 2020; Figure 3). However, although administration of D1/D5 antagonist significantly impacted locomotion (see Supplementary Figure 1B), the general amount of movement or head direction during the exposure sessions were not correlated with the subsequent performance of the observers.

The current results do not exclude the participation of other parallel neuromodulatory circuits. When learning from conspecifics, the observer’s attentional and memory resources are focused on the demonstrator’s exploration and performance of skilled behaviors (Hoppit and Laland, 2013). Therefore, it is likely that the cholinergic and noradrenergic systems play prominent roles in the modulation of attention, arousal, memory formation (Hasselmo, 1999; Fitzpatrick et al., 2019). Moreover, social interactions during the exposure sessions likely engage oxytocin and serotonin signaling (Dölen et al., 2013). Ultimately, it will be necessary to assess the relative impact of each mechanism on the neural plasticity mechanisms that support social learning.

## Methods

### Experimental animals

Gerbil (Meriones unguiculatus, n=117) pups were weaned at postnatal day (P) 30 from commercial breeding pairs (Charles River). Littermates were caged together, but separated by sex, and maintained in a 12 h light/dark cycle. All procedures related to the maintenance and use of animals were approved by the University Animal Welfare Committee at New York University, and all experiments were performed in accordance with the relevant guidelines and regulations.

### Behavioral setup

The behavioral setup was similar to Paraouty et al. (2020). Gerbils were placed in a plastic test cage (dimensions: 0.4 × 0.4 × 0.4 m) that was housed in a sound attenuation booth (Industrial Acoustics; internal dimensions: 2.2 × 2 × 2 m), and observed via a closed-circuit monitor. Auditory stimuli were delivered from a calibrated free-field tweeter (DX25TG0504; Vifa) positioned 1 m above the test cage. Sound calibration measurements were made with a ¼ inch free-field condenser recording microphone (Bruel & Kjaer). A pellet dispenser (Med Associates Inc, 20 mg) was connected to a food tray placed within the test cage, and a nose port was placed on the opposite side. The nose port and food tray were equipped with IR emitters and sensors (Digi-Key Electronics; Emitter: 940 nm, 1.2V, 50 mA; Sensor: Photodiode 935 nm 5 nS). Stimuli, food reward delivery, and behavioral data acquisition were controlled by a personal computer through custom MATLAB scripts and an RZ6 multifunction processor (Tucker-Davis Technologies).

### Stimuli

For the sound discrimination task, the Go stimulus consisted of amplitude modulated (AM) frozen broadband noise tokens (25 dB roll-off at 3.5 kHz and 20 kHz) with a modulation rate of 12 Hz and a modulation depth of 100%. The Nogo stimulus was similar to the Go stimulus, except for the modulation rate which was 4 Hz. Both Go and Nogo stimuli had a 200 ms onset ramp, followed by an unmodulated period of 200 ms which then transitioned to an AM stimuli. The sound level used was 55 dB SPL.

### Video recordings

Videos of the test cage were captured with a Logitech c270-HD webcam (30 frames per second, Best Buy). We used the open source software: DeepLabCut (Mathis et al., 2018) on a Windows 10 machine (Dell Precision 5820, 64-bit operating system) to track the gerbil’s position in the test cage during practice sessions. The network was trained with 1,030,000 iterations using a total of 1064 labeled frames (labeling of nose, left ear, right ear, and tail base) and tested on a set of 200 frames. Manual evaluation of labeling accuracy was achieved by comparing the labels acquired from the network on the test set with the manual labels.

### Sound recordings

Audio recordings were captured with two Dodotronic microphones (Ultramic 384K_BLE), placed on either side of the test cage. We used the open source software: DeepSqueak (Coffey et al., 2019) on a Windows 10 machine (Dell Precision 5820, 64-bit operating system) to track the vocalizations of both the demonstrator animal and the observer animal in each session. DeepSqueak-screener was used to first create a library of gerbil calls. A detection network was then produced using this library of gerbil vocalizations. All audio recordings were analyzed using this custom gerbil detection network, followed by a post-hoc denoising stage and manual confirmation of all individual calls. Due to the physical barrier (i.e., the divider) separating the observer and demonstrator animal, only a subset of calls were caught by both speakers. Thus, the majority of calls (85% on average across all sessions) of the demonstrator was only caught by the demonstrator’s microphone and the majority of calls (83% on average across all sessions) of the observer animal was only caught by the observer’s microphone. For the subset of calls that were caught by both microphones, the calls were attributed to the side with the highest power (in dB).

### Experimenter trained demonstrator gerbils

Demonstrator gerbils (n=43) were trained by the experimenters on a sound discrimination task. The demonstrators were placed on controlled food access prior to the start of training, and all animals were trained using an appetitive reinforcement operant conditioning procedure (see Paraouty et al., 2020). Animals first learned to approach the food tray and receive food pellets (Bio Serv) when the Go stimulus (12 Hz AM noise) was played. Animals were then trained to reliably initiate Go trials independently by placing their nose in the port. Once animals were performing a minimum of 80 Go trials with a hit rate > 80%, Nogo trials were introduced. The probability of Nogo trials was kept at 30% in order to keep the animal motivated to perform the task. Nogo trials were paired with a 4-second time-out during which the house lights were extinguished and the animal could not initiate a new trial. The presentation of Go and Nogo trials were randomized to avoid animals developing a predictive strategy. For more details on the experimenter training procedure, refer to Paraouty et al. (2020).

Responses were scored as a hit when animals approached the food tray to obtain a food reward upon Go trials. If animals re-poked or did not respond during the 5-second time window following a Go stimulus, it was scored a miss. During Nogo trials, responses were scored as a false alarm when animals incorrectly approached the food tray. If animals re-poked or did not respond during the 5-second time window following a Nogo stimulus, then it was scored a correct reject. Hit and FA rates were constrained to floor (0.05) and ceiling (0.95) values to avoid d’ values that approach infinity. A performance metric, d prime (d’) was then calculated for each session by performing a z-transform of both Hit rate and False Alarm values: d’ = z(Hit rate) – z(False Alarm rate) (Green and Swets, 1966). To qualify as a demonstrator, animals were required to perform the task with a d’ > 1.5.

### Social learning paradigms

During each *exposure session* (see Video 1), a demonstrator gerbil performed the discrimination task in the presence of a naïve, untrained observer gerbil (n=74, age=96.9 ± 32.7). No animals were excluded from the study. As shown in Paraouty et al., 2020 (Supplementary Figure 3), we found no significant difference between observers that were paired with either same-sex cagemates or same-sex non-cagemates. Thus, in the current study, we used both cagemates and non-cagemates demonstrators. In addition, the age difference between observer and demonstrator was identical for cagemates, or slightly older demonstrators for non-cagemates. As in Paraouty et al., 2020 (Supplementary Figure 5), we did not observe a significant correlation between the demonstrator’s performance nor age and the observer’s subsequent performance on the task.

Both the demonstrator and the naïve observer gerbil were placed on controlled food access. A divider (acrylic sheet) was placed within the test cage to separate the demonstrator compartment from the observation or exposure compartment (see Paraouty et al., 2020). The naïve observer animal could thus see the demonstrator perform the task and was also exposed to the Go and Nogo sound stimuli as the speaker was located 1 m above the test cage. For the observers, a nose port and food tray were present only on the demonstrator’s compartment, allowing the demonstrator to initiate and perform trials (Figure 1A, left). For the yoked reward observers, a food tray was also present in the observer’s compartment, and the food reward of the demonstrator animal was yoked to that of the observer (Figure 1D, left). In both conditions, the naïve observers were exposed to a minimum of 80 Go trials and 20 Nogo trials performed by the demonstrator gerbils in each exposure session.

After five daily exposure sessions, the divider was removed after the final day of exposure, and the naïve observer gerbil was then allowed to practice the task (*practice session*). The observer’s food tray (when present, i.e., for the yoked reward conditions) was also removed during the practice sessions. During the first and second practice sessions, the observer gerbil was given the benefit of no more than five experimenter-triggered Go trials. These experimenter-triggered Go trials were initiated only when an animal was touching the nose port. This method of manually initiating Go trials was identical to the one used to train demonstrators, in order to maximize the animal’s interest in the nose port object. Except for these experimenter-triggered Go trials, all Go trials were initiated by the gerbil. In order to limit a source of high variance, we only introduced Nogo trials once an observer animal was reliably initiating Go trials and performed >25 Hits. False alarm trials were paired with a 2-second time-out on the second day of Nogo trial introduction. For all following practice days, a 4-second time-out was used when animals false alarmed. A d’ was computed for all practice sessions during which a minimum number of 15 Nogo trials were presented (see Paraouty et al., 2020). This was the standard procedure used for all observer groups tested here.

### Drug and saline injections

Prior to each exposure session, a subset of naïve observer gerbils was briefly anaesthetized with isoflurane and given an intra-peritoneal injection of either physiological saline (0.9% NaCl, n=13) or a D1/D5 dopamine receptor antagonist, SCH-23390 (0.025 mg/kg, volume=0.06ml, n=10) or a D1/D5 dopamine receptor agonist, SFK-38393 (5.0 mg/kg, volume=0.1 ml, n=11). In addition, a subset of yoked reward observer gerbils was injected with physiological saline (0.9% NaCl, n=6) or the D1/D5 dopamine receptor antagonist, SCH-23390 (0.025 mg/kg, volume=0.06ml, n=6). The doses were initially based from previous studies in gerbils (Barnes et al., 2016 ; Schicknick et al., 2008; 2012). Following injections, animals were allowed to fully recover in a recovery cage (for 15-25 minutes) before the exposure session began. Higher doses of both the agonist and antagonist produced noted behavioral and motor effects (excessive grooming and hyperactivity for the agonist, and reduced motor behavior for the antagonist).

### Performance measures and statistical analyses

Due to limited litter sizes, a given litter could not be split into all conditions. However, animals from one given litter was used for at least 2 conditions. In addition, for all groups, data was collected from at least 3 different litters in order to avoid any litter-specific biases. No litter differences were observed during the experimenter-training stages of the demonstrator animals. A performance measure (d’) was calculated for each animal: d’ = z(Hit rate) – z(False Alarm rate). Hit and False Alarm rates were constrained to floor (0.05) and ceiling (0.95) values. To avoid high variance, d’ was only computed for sessions in which the observer gerbil performed > 15 Nogo trials. For the computation of the mean d’ line, we used all values of d’ and attributed a zero to all NaN values of d’ (i.e., when an animal was initiating < 15 Nogo trials, hence no d’ value could be computed). However, for all statistical tests and for the computation of the mean number of days taken by each experimental and control group to reach a criterion d’ of 1.5, only actual d’ values were used. We first checked whether the number of days for each group to reach a criterion performance d’ of 1.5 were normally distributed using the Shapiro Wilk test of normality. As the latter were not sufficiently Gaussian, we chose to perform non-parametric tests. All group level statistical tests and effect size calculations were performed using JMP Pro 14.0 on a Mac platform. Comparisons were carried out using the Wilcoxon rank sum test or the Steel Dwass comparisons, as indicated. For comparisons of all groups tested, one-way ANOVAs were computed followed by post-hoc multiple comparisons analyses, and alpha values were Holm-Bonferroni-corrected.

## Abbreviation

AM: amplitude modulation

## Acknowledgements

The work was supported by the Fyssen Foundation (NP) and R01DC011284 (DHS).

## Video legend

**Video 1**

Social learning of an auditory discrimination task. A naïve *observer* gerbil was permitted to experience a trained conspecific *demonstrator* perform an auditory Go-Nogo discrimination task. The naïve observer gerbil was exposed to a minimum of 80 Go trials and 20 Nogo trials in each exposure session. The performance of the demonstrator during those 5 exposure sessions is shown in Figure 1 (brown lines). Following five such exposure sessions, the naive observer gerbil was subsequently permitted to practice the task. The performance of the observer gerbil during the subsequent practice sessions is shown in Figure 1 (black lines). The markings on top of the demonstrator and the observer gerbil were obtained using DeepLabCut with 4 labelled features marked: nose tip, left ear, right ear, and tail base.

## Supplementary Information

**Supplementary Figure 1**

***Administration of dopamine antagonist impacts the general behavior of observers*. A.** Illustration of the distance travelled by a demonstrator and an observer during 1 example exposure session. **B.** Mean distance explored per exposure day for a subset of 5 observers in each of the different observer groups: left- naïve observers (black) and yoked reward observers (orange); middle- observers which received saline (blue), or a dopamine antagonist (red), or a dopamine agonist (green); right- yoked reward observers which received saline (blue), or a dopamine antagonist (red). The analysis of distance travelled was performed on the same number of frames (first 20 minutes) of each of the five daily exposure sessions. **C.** Illustration of the head orientation of a demonstrator and an observer during 1 example exposure session. **D.** Mean ratio of head orientation for the 0-18° which corresponds to the food tray for the same subset of 5 observers (as in B). Asterisks denote statistically significant post-hoc differences at the following levels: * *p*<0.05 and \*\**p*<0.01.

**Supplementary Figure 2**

***D1/D5 antagonist injections during exposure negatively impacts subsequent performance.* A.** Mean total number of trials (± SEM) initiated by all observers in the seven different experimental groups of observers across practice days (Go and Nogo trials). **B.** Mean hit rate (± SEM) of all observers across practice days. A mixed-model ANOVA revealed significant group differences, and post-hoc comparisons confirmed the significantly poorer Hit rates of the D1/D5 antagonist observer groups as compared to all other groups (p<0.001) during the early days of practice. No significant group difference was found between the other observer groups.

**Supplementary Figure 3**

***Summary Table of the differential experimental paradigms, described in the current study and in Paraouty et al. (2020).*** Illustration of exposure sessions for all conditions (left).

